# Bridging Transcriptome Regulation and Cellular Operational Efficiency with the Locality and Caching System Optimization Principles

**DOI:** 10.1101/2022.01.11.475967

**Authors:** Wen Jiang, Denis Feliers, W. Jim Zheng, Fangyuan Zhang, Degeng Wang

## Abstract

Gene expression is time-consuming and sequentially more so from bacteria to yeast to human, rendering human cells vulnerable to proteomic-response and operational latency. Computers once suffered such latency, imposed by much-slower information retrieval (hard-drive (HD) to memory to central-processing-unit (CPU)) than CPU execution. Optimization principles, namely, spatiotemporal-locality-principles that control specialized user-programs and caching that controls operating system (OS) kernel (the HD-CPU information flow channel), successfully mitigated the latency by gearing the memory towards near-future or high-priority CPU needs. We report evidence that the principles similarly act in cellular latency-mitigation via analogizing genome-mRNA-protein gene-expression to HD-memory-CPU information-retrieval, and transcriptome to memory. First, temporal-locality-principle is equivalent to mRNA stabilization-by-translation regulation and controls specialized cellular functions. Second, caching is equivalent to cytoplasmic mRNA sequestration. Highly sequestered mRNAs defy the locality-principle. In both cells and computers, caching controls the information channels; gene expression machinery and their regulators, *i.e*., the cellular channel (OS-kernel equivalent) that regulates arguably all cellular processes, are top sequestered mRNAs. Third, mRNA-caching contributes to the mRNA-protein expression discrepancy. Thus, locality and caching principles control specialized and core cellular functions, respectively, orchestrating transcriptome regulation and bridging it to cellular operational efficiency.

## Introduction

Gene expression is one of, if not the, most fundamental cellular processes. It goes hand-in-hand with the control of essentially all biochemical and regulatory pathways, activating and inhibiting a pathway by increasing and decreasing the abundance of corresponding proteins, respectively. On the other hand, the multi-stepped process is time consuming, and sequentially more so from bacteria to yeast and to mammalians; for instance, transcription elongation rate is 42-54 nucleotides per second in bacteria, but only 22-25 nucleotides per second in human. This prolonged delay from transcription activation to protein production in human cells would, if not mitigated, render it impossible for the cells to make rapid responses to environmental signals – a severe system latency issue. On the other hand, the gene expression process consumes ATP and GTP as energy sources, and nucleotides and amino acids as building blocks, levying significant metabolic overhead on the cells. Thus, gene expression regulation is a critical cellular operational optimization problem; human cells must have evolved elaborate regulatory strategies to strike the optimal balance, *i.e*., bypassing the latency and achieving operational efficiency without incurring excessive wasteful metabolic expenditure.

At the transcriptome level, extensive studies have characterized many regulatory mechanisms and revealed tremendous complexity; for instance, the binding site mapping for hundreds of RNA binding proteins (RBP) and the identification of many non-coding RNAs (1,2). MicroRNA (miRNA), perhaps the most studied non-coding RNAs, controls >60% of mRNAs (3–5). While transcription regulation is the primary controller of mRNA production, these posttranscriptional regulators control mRNA translation and/or degradation via cognate binding sites, which are mostly in mRNA un-translated regions (UTR), especially 3’-UTR. The prevalence of these regulatory mechanisms is underscored by the observation that UTR occupies, on average, 50% of the length of a mRNA (6); in case of the CREB1 gene, it is 90%. These regulatory mechanisms lead to gene expression complexity, often manifested as discrepancy among key gene expression parameters, namely, the transcription rate, mRNA abundance, mRNA translation activity and protein abundance (7–16). This was noticed prior to the genomic era (17,18), confirmed by the first simultaneous genome-wide analysis of multiple gene expression parameters (19), and then quickly by numerous such studies (20–28); for instance, discrepancy between mRNA and protein abundance has emerged as a prominent recurring theme in latest proteogenomic studies (29–31).

However, the regulatory mechanisms and the complexity are not studied in close conjunction with cellular operational efficiency. That is, though repeatedly observing the WHAT, we do not yet know the underlying WHY and HOW. The operational advantages these regulatory mechanisms confer to the cells remain unclear, and we do not understand well the functional and mechanistic underpinning for the discrepancy among key gene expression parameters. Additionally, while biochemical details of many regulatory mechanisms are known, we have yet to uncover the underlying principles for cellular orchestration of multiple regulatory mechanisms among the genes and functional groups. It is more likely that multiple mechanisms act in concert with one another, than individual mechanisms acting alone, to support efficient execution of cellular processes and operations.

Thus, we strived to evaluate transcriptome regulation in the context of cellular needs to mitigate the key operation efficiency issue – the latency at the mRNA level, *i.e*., the lengthy time delay from transcription to mRNA translation during the gene expression process, in human cells. To mitigate the latency, the cells need to orchestrate transcription, mRNA degradation and mRNA translation activities among all the genes. There are two seemingly contradictory objectives – expression of mRNAs in advance of cellular needs for their proteins and minimization of wasteful expression of mRNAs that are degraded without ever being translated. Additionally, selective degradation of existing mRNAs occurs constantly, conceivably to free up resources to accommodate incoming nascent mRNAs; ideally, only those mRNAs that are neither currently translating nor expected to be so in the near future are subject to degradation. Unfortunately, we do not have a cohesive model for how the cells achieve these objectives.

This latency issue, on the other hand, is not unique to the cells. The computer information retrieval process from hard drive (HD) to memory, and then to central processing unit (CPU) cache, is strikingly similar to the gene expression process (16,32,33). They both function to support system operation through dynamic retrieval of permanently stored digital information–the binary information in the HD in computers and the quadruple (A, T, C and G) genetic information in the genomes in cells; and both are via an intermediate step–the memory in computers and the mRNA level in the cells. Similar to the tardiness of gene expression in human cells, the HD-to-CPU information retrieval is much slower than CPU execution cycle; the CPU might stay idle for extended periods of time while waiting for information for the next cycle–a latency issue once severely plaguing the computing community (34). Moreover, for core system functions, such delay would destabilize and potentially crash the whole system. Memory management/optimization principles, namely, spatiotemporal locality and caching, were developed; the goal is efficient speculative loading and purging of information, dedicating the limited memory capacity to current and near future CPU tasks as well as to the maintenance of system stability (34,35). Implementation of these principles successfully mitigated the latency issue. Consequently, that these memory management and optimization principles also apply to transcriptome regulation becomes an interesting, and powerful, hypothesis.

To test this hypothesis, we performed a multi-omics comparative study of gene expression to the HD-to-CPU information flow. Excitingly, our results support the hypothesis. First, the gene expression process becomes sequentially more selective, *i.e*., focusing on sequentially less genes, from transcription to translation; the trend is similar to the sequential HD-to-CPU capacity decreases, and thus selectivity increases, in computers. Second, the temporal locality principle that governs speculative memory information purging in computers is equivalent to the mRNA stablewhen-translating regulatory mechanism, and controls many cellular functions, as exemplified by major metabolic pathways. Third, many core functions, as exemplified by basal transcription and translation machineries, defy the stable-when-translating regulatory mechanism. Instead, high levels of mRNA cytoplasmic sequestration, *i.e*., stable but not translating, were observed. Strikingly, these functions and the permanently cached functions in computers are equivalent to each other, acting as the information flow channel in the respective system. That is, permanent memory caching is equivalent to mRNA sequestration. Fourth, mRNA sequestration correlates well with mRNA-protein expression discrepancy independently observed in a proteogenomic study. In short, coordination of the locality and caching principles, and their centrality to computer operation optimization, outlined cellular orchestration of multiple transcriptome regulatory mechanisms in the context of operational latency mitigation, closing the conceptual gap to bridge transcriptome regulation and cellular operation efficiency.

## Materials and Methods

### Tissue Culture and mRNA Isolation for RNA-seq Analysis

As previously described (36–38), the human HCT116 cells were cultured in a serum-free medium (McCoy’s 5A (Sigma) with pyruvate, vitamins, amino acids and antibiotics) supplemented with 10 ng/ml epidermal growth factor, 20 μg/ml insulin and 4 μg/ml transferrin. Cells were maintained at 37 °C in a humidified incubator with 5% CO_2_.

To extract mRNA for RNA-seq analysis, RNeasy kit (Qiagen) was used to extract total RNA from the HCT116 cells according to manufacturer’s specification. GeneRead Pure mRNA Kit (Qiagen) was then used to isolate mRNA from the total RNA for Illumina NGS sequencing according to manufacturer’s specification.

### GRO-seq Analysis

Global run-on was done as previously described (6,39–42). Briefly, two 100 cm plates of HCT116 cells were washed 3 times with cold PBS buffer. Cells were then swelled in swelling buffer (10 mM Tris-pH 7.5, 2 mM MgCl2, 3 mM CaCl2) for 5 min on ice. Harvested cells were re-suspended in 1 ml of the lysis buffer (swelling buffer with 0.5% IGEPAL and 10% glycerol) with gentle vortex and brought to 10 ml with the same buffer for nuclei extraction. Nuclei were washed with 10 ml of lysis buffer and re-suspended in 1 ml of freezing buffer (50 mM Tris-pH 8.3, 40% glycerol, 5 mM MgCl2, 0.1 mM EDTA), pelleted down again, and finally re-suspended in 100 μl of freezing buffer.

For the nuclear run-on step, re-suspended nuclei were mixed with an equal volume of reaction buffer (10 mM Tris-pH 8.0, 5 mM MgCl2, 1 mM DTT, 300 mM KCl, 20 units of SUPERase-In, 1% Sarkosyl, 500 μM ATP, GTP, and Br-UTP, 2 μM CTP) and incubated for 5 min at 30 °C. Nuclei RNA was extracted with TRIzol LS reagent (Invitrogen) following manufacturer’s instructions, and was resuspended in 20 μl of DEPC-water. RNA was then purified through a p-30 RNAse-free spin column (BioRad), according to the manufacturer’s instructions and treated with 6.7 μl of DNase buffer and 10 μl of RQ1 RNase-free DNase (Promega), purified again through a p-30 column. A volume of 8.5 μl 10 × antarctic phosphatase buffer, 1 μl of SUPERase-In, and 5 μl of antarctic phosphatase was added to the run-on RNA and treated for 1 hr at 37 °C. Before proceeding to immuno-purification, RNA was heated to 65 °C for 5 min and kept on ice.

Anti-BrdU argarose beads (Santa Cruz Biotech) were blocked in blocking buffer (0.5 × SSPE, 1 mM EDTA, 0.05% Tween-20, 0.1% PVP, and 1 mg/ml BSA) for 1 hr at 4 °C. Heated run-on RNA (~85 μl) was added to 60 μl beads in 500 μl binding buffer (0.5 × SSPE, 1 mM EDTA, 0.05% Tween-20) and allowed to bind for 1 hr at 4 °C with rotation. After binding, beads were washed once in low salt buffer (0.2 × SSPE, 1 mM EDTA, 0.05% Tween-20), twice in high salt buffer (0.5% SSPE, 1 mM EDTA, 0.05% Tween-20, 150 mM NaCl), and twice in TET buffer (TE pH 7.4, 0.05% Tween-20). BrdU-incorporated RNA was eluted with 4 × 125 μl elution buffer (20 mM DTT, 300 mM NaCl, 5 mM Tris-pH 7.5, 1 mM EDTA, and 0.1% SDS). RNA was then extracted with acidic phenol/chloroform once, chloroform once and precipitated with ethanol overnight. The precipitated RNA was re-suspended in 50 μl reaction (45 μl of DEPC water, 5.2 μl of T4 PNK buffer, 1 μl of SUPERase_In and 1 μl of T4 PNK (NEB)) and incubated at 37 °C for 1 hr. The RNA was extracted and precipitated again as above before being processed for Illumina NGS sequencing.

### Fragmentation of cytoplasmic extract and mRNA extraction

Cytoplasmic extract fragmentation was performed as previously described (43,44). Briefly, the HCT116 cells were incubated with 100μg/ml cycloheximide for 15 minutes, washed three times with PBS, scraped off into PBS, and then pelleted by micro-centrifugation. Cell pellet was homogenized in a hypertonic re-suspension buffer (10 mM Tris (pH 7.5), 250 mM KCl, 2 mM MgCl2 and 0.5% Triton X100) with RNAsin RNAse inhibitor and a protease cocktail. Homogenates were centrifuged for 10 min at 12,000 g to pellet the nuclei. The post-nuclear supernatants were laid on top of a 10-50% (w/v) sucrose gradient, followed by centrifugation for 90 min at 200,000 g. The polysomal and non-polysomal fractions were identified by OD254 and collected. RNeasy kit (Qiagen) was used to extract RNA from both fractions according to manufacture’s specification. GeneRead Pure mRNA Kit (Qiagen) was then used to isolate mRNA for Illumina NGS sequencing from the RNA according to manufacture’s specification.

### Illumina NGS Sequencing

Sequencing libraries were generated with the Illumina TruSeq RNA Sample Preparation Kit. Briefly, RNA molecules were fragmented into small pieces using divalent cations under elevated temperature. The cleaved RNA fragments are copied into first strand cDNA synthesis using reverse transcriptase and random primers. This was followed by second strand cDNA synthesis using DNA Polymerase I and RNase H. These cDNA fragments were end-repaired using T4 DNA polymerase, Klenow polymerase and T4 polynucleotide kinase. The resulting blunt-ended fragments were A-tailed using a 3’–5’ exonuclease-deficient Klenow fragment and ligated to Illumina adaptor oligonucleotides in a ‘TA’ ligation. The ligation mixture was further size-selected by AMPure beads and enriched by PCR amplification following Illumina TruSeq DNA Sample Preparation protocol. The resulting library is attached and amplified on a flow-cell by cBot Cluster Generation System.

The sequencing was done with an Illumina HiSeq 2000 sequencer. Multiplexing was used to pool 4 samples into one sequencing lane. After each sequencing run, the raw reads were pro-processed to filter out low quality reads and to remove the multiplexing barcode sequences. The GRO-Seq, RNA-seq and polysome-seq datasets are available through the NCBI GEO database (accession number GSE111222). The non-polysome-Seq data was submitted to the GEO database (accession number GSE196299).

### NGS Data Analysis

The sequencing reads were mapped to the UCSC hg19 human genome sequences with the STAR software (45), using the default input parameter values. For each sample, at least 80% of the reads were successfully mapped. For the sake of consistency across the four transcriptome regulation parameters, we counted the reads for each gene for the exon regions only. The counting was performed with the HTSeq-count software (46), and the counts were then transformed into Reads Per Kilo-base Per Million Mapped Reads (RPKM) values. 12921 genes have a minimal RPKM value of 1 for at least one of the three parameters, and were considered expressed in the HCT116 cells. Linear regression of log-transformed data was used to examine consistence between biological replicate samples.

### The Kyoto Encyclopedia of Genes and Genomes (KEGG) and Gene Ontology (GO) gene sets

The KEGG and GO functional gene sets were downloaded from the Molecular Signatures Database (MSigDB) version 7.4 at the Gene Set Enrichment Analysis (GSEA) website (https://www.gsea-msigdb.org/gsea/index.jsp) (47,48). The KEGG gene set contains 186 gene sets. The GO molecular function (MF) set contains 1708 sets, and the biological process (BP) and the cellular component (CC) sets 7481 and 996 sets, respectively.

### Statistical Analysis

The R open source statistical software (version 3.6.2) installed on a Mac Pro desktop computer was used for statistical analysis. Outlier identification, student t-test, descriptive statistical parameter calculation, correlation coefficient calculation, linear regression, loess regression and other statistical procedure are all done with this R software.

### Statistics of pair-wise sequestration index analysis

We calculated the pair-wise differences in the sequestration indexes (sequ^diff^) among all the genes. For a set of gene pairs, the seq^diff^ parameter should follow a normal distribution with a mean of 0 and a variance (var) given by the following equation:

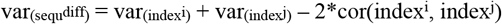

For the whole set of gene pairs, the following steps are used to calculate the var and standard deviation (SD):

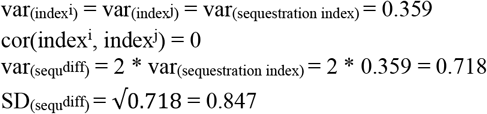

### GO Similarity Analysis

Pairwise GO similarity scores between human genes were computed as previously described (49–51). Briefly, for each gene, we first generated GO fingerprint – a set of ontology terms enriched in the PubMed abstracts linked to the gene, along with the adjusted p-value reflecting the degree of enrichment of each term. The GO similarity score quantifies similarity between the GO fingerprints of corresponding gene pair. For detail about GO fingerprint generation and similarity calculation, please see description in previous publications (50).

## Results

### Gene expression and information retrieval in computers as instances of the information theory in support of system operation

It is long observed that the cells and the computers share many common architectural and operational features, though the former rely on controlled biochemical processes, *i.e*., the biochemical reactions/flow in the biochemical/regulatory network, and the latter on controlled electrical flow (52–55). We have previously reported the similarity between gene expression and the HD-to-CPU information retrieval and their centrality to respective system operations (16,32,33). In this study, we unified the two as respective implementation of the components of the Shannon information theory: information encoding/storage, the execution level where the information is decoded into actions, as well as the channel through which the information is dynamically retrieved from the storage level to the execution/decoding level (56) (Figs. 1 A and B). In the cells, the quadruple genetic (A, T, C and G) information is stored in the chromosomes; in computers, it is the binary information stored in the hard drive (HD). At the execution/decoding level, the binary information retrieved into the CPU cache is decoded into corresponding electrical flow patterns in the CPU to execute specific Boolean algebraic transformation in computers; in the cells, mRNAs form polysome complex with ribosomes, translating the quadruple information into proteins with specific 3-dimensional structures and functional domains to carry out corresponding biochemical functions – often to catalyze a biochemical reaction/transformation.

**Figure 1.**
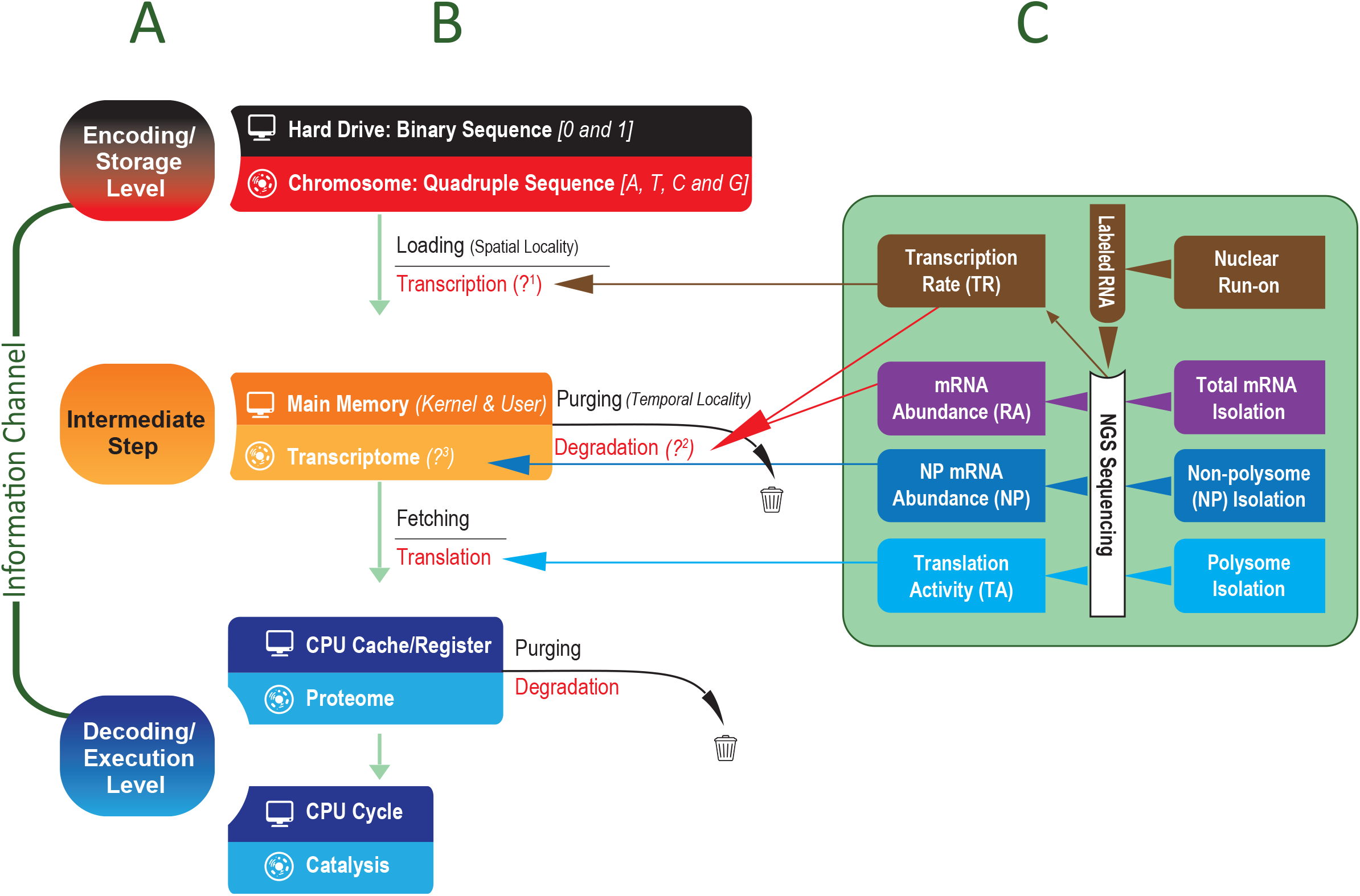
Analogizing the gene expression process to the computer information retrieval process via the Shannon information theory (A & B), and our strategy for experimental exploration of the analogy (C). **A:** outline of the three components of the information theory (encoding/storage, information-channel, and decoding/execution), with the information channel enclosing an intermediate step. **B:** Comparing cellular and computer instances of the theory: harddrive (HD) binary *vs* genomic sequences; memory *vs* mRNA; and CPU cycles *vs* protein biochemical actions. Parenthesized italic texts at each information-retrieval step specify the computer optimization principles, and parenthesized question marks (?) ask whether equivalent principles act in the cells. The sequentially narrowing textboxes denote HD-to-CPU capacity decrease in computer. **C:** Our experimental strategy to answer the questions. Log-phase HCT116 cells were split into multiple aliquots. One aliquot was used for RNA-seq analysis to measure steady-state mRNA abundance (RA) (purple textboxes and arrows). One aliquot was used for GRO-seq analysis to measure transcription rate (TR) (brown textboxes, lines and arrows). Another aliquot was used to isolate and quantify both polysome-associated mRNA/translation activity (TA) (light blue textboxes and arrows) and non-polysome-associated cytoplasmic mRNA (NP) (blue textboxes and arrows). The arrowed lines between panels B and C map experimentally measured parameters to the targeted gene expression steps.

Intriguingly, the information channel encompasses a middle step between the storage and the execution/decoding levels in both systems – the mRNA step in the cells and the main memory in computers. In the cells, the molecular biology central dogma outlines the genome-to-proteome information flow via the middle step, and we have argued that the gene expression machineries and their regulators constitute the information channel (33). Briefly, the basic gene expression machineries, such as RNA polymerase II and ribosome, perform the task; cellular signaling, on the other hand, contributes to the specificity, *i.e*., which genes are transcribed and which mRNAs are translated at any particular moment, by controlling cognate transcription factors and translation regulators.

This prompted us to closely examine the HD-to-CPU information flow via the main memory to identify the information channel in computers, leading our attention to the operating system (OS) kernel. One feature of the process is sequentially decreasing capacity, and thus sequentially increasing information content selectivity, from HD to memory and to CPU cache (Fig. 1B). There is a constant need to purge information already in the memory to free up space for the incoming information. Additionally, the information stored in the HD are organized into two categories: specialized functions commonly termed user programs as exemplified by Microsoft Word; and core system functions commonly known as OS kernel that maintains system stability and running environments for all user programs (Fig. 1B). The kernel is described in the literature as managing the hardware resource, *i.e*., dynamically and optimally allocating the resource among competing user programs, which, as discussed below, can be re-phrased as collectively carrying out the dynamic HD-memory-CPU information retrieval process. Memory management is one major kernel function; the other functions are the file system and process control. The file system implements efficient HD information organization and keeps track of the locations of individual pieces of information, enabling quick retrieval of specified information into the memory. Memory management controls both information loading into, and purging out of, the memory. The kernel’s process-control function controls the information flow from memory to CPU cache and CPU execution, which is equivalent to the translation step and catalysis of biochemical reactions by the produced proteins. Thus, it is conceivable that the OS kernel acts as the channel for the dynamic HD-to-CPU information flow in the computer, *i.e*., the equivalent to gene expression machinery and their regulators.

Intrigued by this potential functional equivalence, we embarked on an experimental investigation, aiming to determine whether and how much the cell-computer analogy sheds insight into human cell gene expression complexity and operational efficiency.

### The system latency issue and its mitigation in computers

As mentioned earlier, in both systems, the retrieval process imposes potential latency due to the inevitable time delay from the storage level to the execution level. Though aware that the delay becomes sequentially longer from bacteria to yeast and to humans, we do not yet systematically understand how human cells overcome the latency.

However, we fully understand this latency and its mitigation in computers. The HD-to-memory retrieval is much slower than that from memory to CPU cache, which is, in turn, much slower than a CPU decoding/execution cycle (34). As discussed below, the latency was mitigated with key memory management and optimization principles that predict near-future and high-priority CPU needs, enabling speculative information loading into, and purging out of, the memory.

Briefly, though necessitated by the latency, the speculation is not free of side-effects and demands elaborate control. That is, speculation inaccuracy is detrimental, but certain to occur. For loading, a portion of the speculatively loaded information might get purged without ever being requested by the execution step, wasting memory space and the system resource dedicated to the loading operation. Similarly, a side effect of speculative purging is unfortunate removal of soon-to-be-needed information and having to quickly reload it, a situation termed “thrashing” that is even more wasteful and detrimental than a speculative loading error. In general, higher levels of speculation, *e.g*., a larger volume of loaded information per loading operation, lead to higher levels of speculation inaccuracy.

To maximize system operational efficiency, computer memory management seeks optimal tradeoff among the needs for speculative information loading into and purging out of the memory, incurrence of counterproductive speculation inaccuracy and other system operation parameters. For instance, for user programs, this is achieved via OS-kernel implementation of spatial and temporal locality principles (Fig. 1B); the principles, to be discussed in detail later, bestow predictive power for near-future CPU cycles, reducing counterproductive speculation inaccuracy and maximizing the likelihood that information for near future CPU cycles is always in the memory. That is, the memory is dynamically geared towards current and near-future CPU needs to mitigate latency in user program execution (34).

### Extending the cell-to-computer analogy and a multi-omics experimental testing strategy

We asked whether and how much the computer-cell analogy can be extended to better understand latency-mitigation and gene expression complexity in human cells. Specifically, we aimed to test three hypotheses: 1) the gene expression process shares the sequential enhancement of selectivity from storage/encoding to execution/decoding steps; 2) principles implemented in computer OS kernel for memory management also act in transcriptome regulation to mitigate cellular operational latency, as denoted by the question marks (?^1^ and ?^2^) in figure 1B (34); and 3) organization of the functions into core system functions and specialized functions would help to understand cellular orchestration of the genes and their functional groups via differential regulatory mechanisms (Fig. 1B, ?^3^).

To answer these questions, we performed multi-omics experimental exploration, *i.e*., simultaneous genome-wide measurement of multiple transcriptome regulation parameters (Figs. 1 B and C). As usual, the RNA-seq was used to measure m<u>R</u>NA <u>a</u>bundance (RA) (Fig. 1C, purple text boxes and arrows). Additionally, we used GRO-seq to measure <u>t</u>ranscription <u>r</u>ate (TR), monitoring transcription as cellular equivalent to computer HD-to-memory information loading (6,39–42) (Figs. 1 B and C, brown text boxes and arrows). Polysome profiling was used to measure <u>t</u>ranslation <u>a</u>ctivity (TA), monitoring translation as the equivalent to information fetching from memory to CPU cache (43,44) (Figs. 1 B and C, light blue text boxes and arrows). And we used TR and RA together, to be described in detail later, to estimate mRNA degradation rate (Figs. 1 B and C, red arrows). The TR, RA, and TA data were previously described (6,42). In this study, we analyzed them in the context of the analogy between gene expression and computer information retrieval. Additionally, the cytoplasmic <u>n</u>on-<u>p</u>olysome-associated mRNA abundance (NP) data was incorporated into this analysis to enable, to be discussed in detail later, a more comprehensive analysis of the analogy (Figs. 1 B and C, blue text boxes and arrows).

### Sequentially higher selectivity from the storage to the execution levels – another architectural similarity to the HD-memory-CPU information-retrieval in computers

As we reported previously, the analysis revealed complexity of transcriptome regulation in the form of discrepancy among the gene expression parameters (Figs. 2, 3 and 4). A trend of sequentially enhanced gene expression selectivity from transcription to translation was observed. Two aspects of the boxplots in figure 2 illustrate the trend schematically. First, the median parameter RPKM values decrease from TR to RA, to NP, and to TA, with all Mann-Whitney-U-test p-values smaller than 1E-16. Second is the sequential TR-RA-NP-TA increases of the variances of the distributions, with F-test p-values all smaller than 1E-16; sequentially more genes are expressed at extreme (high or low) levels. The trend is also shown quantitatively (Table 1). In a word, the cells focus the gene expression resources, *e.g*., the building blocks and the RNA binding proteins, on sequentially less genes as the genetic information flow from the genome to the proteome.

**Figure 2.**
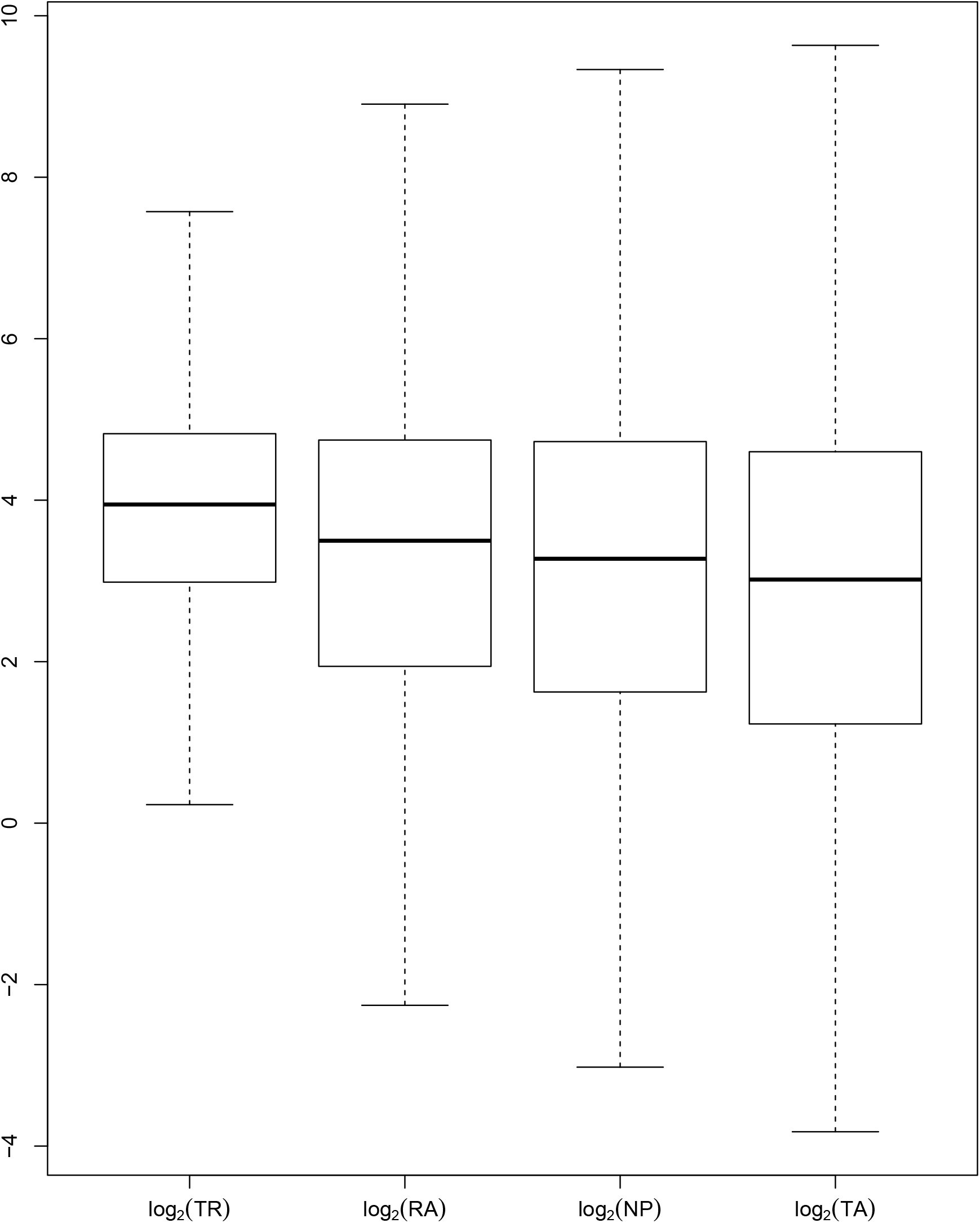
Comparison of log_2_(TR), log_2_(RA), log_2_(NP) and log_2_(TA) distributions. Comparative boxplots of the RPKM values of the four parameters are shown, illustrating the sequentially lower median values, and higher dispersion levels, from TR to TA. The trend is consistent with the HD-to-CPU sequential narrowing of the textboxes in fig. 1B.

**Figure 3:**
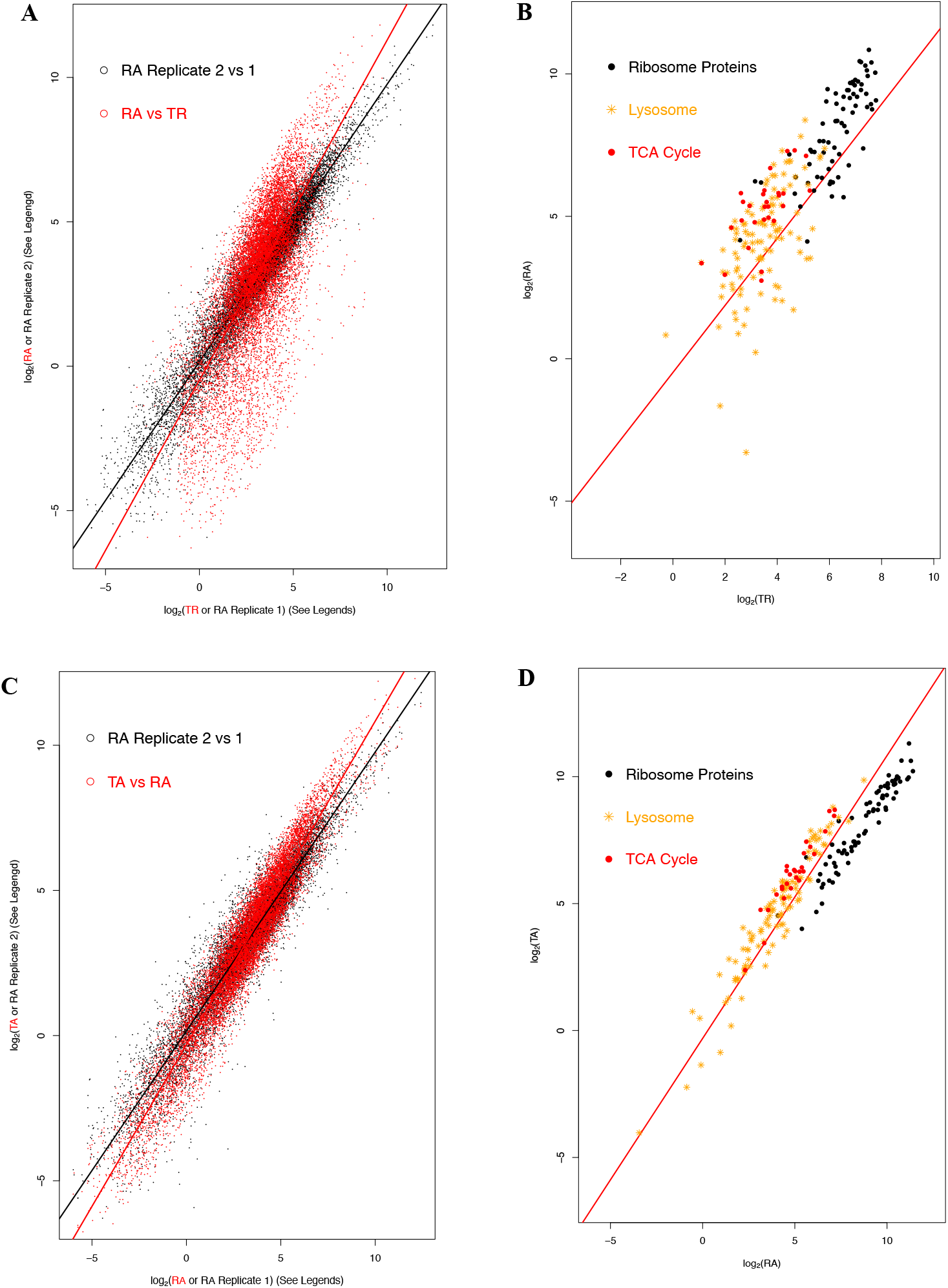
Linear regression analyses to illustrate the discrepancy among TR, RA and TA. **A:** Scatter plot of log_2_(RA) (y-axis) versus log_2_(TR) (x-axis) (red) and log_2_(RA biological replicate 2) (Y-axis) versus log_2_(RA biological replicate 1) (x-axis) (black). Both linear regression lines are also shown. **B:** The log_2_(RA) versus log_2_(TR) scatter plot for three KEGG functional groups: ribosome proteins, lysosome and TCA cycle. The linear regression line (log_2_(RA) versus log_2_(TR)) from A is also shown. **C:** Scatter plot of log_2_(TA) (y-axis) versus log_2_(RA) (x-axis) (red) and of the two RA biological replicates (black). Both linear regression lines are also shown. **D:** The log_2_(TA) versus log_2_(RA) scatter plot for the three functional groups as in **B**: ribosome proteins, lysosome and TCA cycle. The linear regression line (log_2_(TA) versus log_2_(RA)) from C is also shown. The same two RA biological replicates are used in A and C. The patterns in A and C are consistent with the corresponding narrowing of the textboxes in figure 1B. Outliers were removed from the three functional groups.

**Figure 4.**
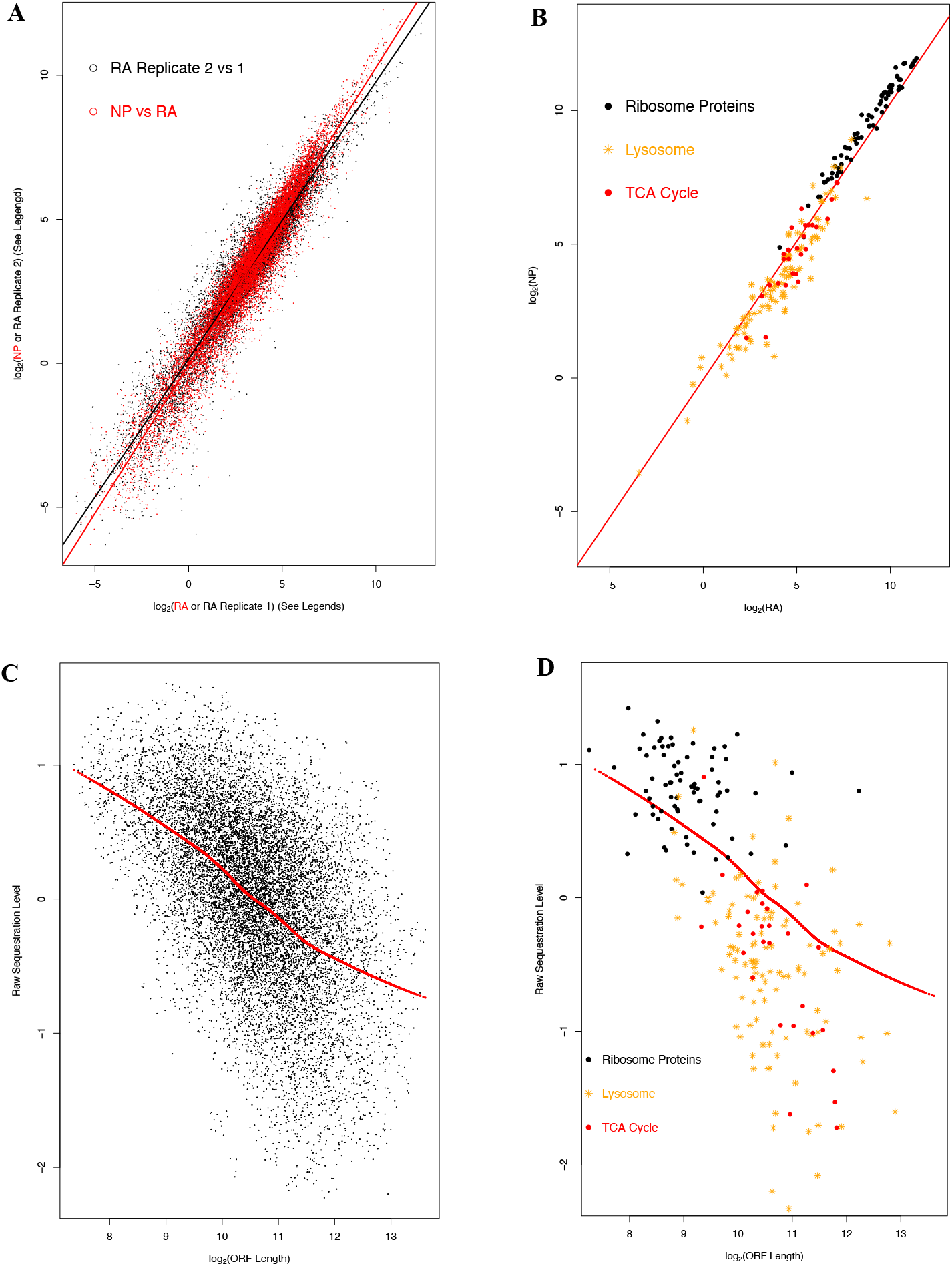
Function-specific pattern of mRNA sequestration and determination of the sequestration index. **A:** Scatter plot of log_2_(NP) (y-axis) versus log_2_(RA) (x-axis) (red) and log_2_(RA biological replicate 2) (Y-axis) versus log_2_(RA biological replicate 1) (x-axis) (black). The two linear regression lines are also shown. The same two RA biological replicates as in figures 3A and 3C are used. **B:** The log_2_(NP) versus log_2_(RA) scatter plot for the three functional groups as in figures 3B and 3D: ribosome proteins, lysosome and TCA cycle. The linear regression line (log_2_(NP) versus log_2_(RA)) from A is also shown. **C**: Scatter plot of the raw mRNA sequestration level versus log_2_(ORF length). The red dots denote the prediction of a loess regression, whose residue is used as the mRNA sequestration index. **D:** Scatter plot (raw mRNA sequestration level versus log_2_(ORF length)) for the three functional groups as in **B**. The loess regression prediction from C is also shown.

**Table 1:** Comparison of statistical features of the profiles of gene expression parameters.

|  | Standard<br>Deviation | Mean | Value Range |  |
| --- | --- | --- | --- | --- |
|  |  |  | From | To |
| $\log_2(\text{TR})$ | 1.53 | 3.85 | -2.68 | 10.41 |
| $\log_2(\text{RA})$ | 2.38 | 3.18 | -6.02 | 12.33 |
| $\log_2(\text{NP})$ | 2.56 | 3.03 | -6.41 | 13 |
| $\log_2(\text{TA})$ | 2.77 | 2.74 | -7.18 | 13.04 |

This trend is, as previously reported (6), also reflected in the slopes in linear regression analyses (Fig. 3). The log_2_(RA) *vs*. log_2_(TR) regression gave a slope of 1.17 – significantly greater than 1 with a smaller than 1E-16 p-value, reflecting that more genes exhibit extreme high or low values in log_2_(RA) than log_2_(TR) (Fig. 3A). Similarly, the log_2_(TA) *vs*. log_2_(RA) regression gave a slope of 1.1 – also significantly greater than 1 with a smaller than 1E-16 p-value, reflecting that more genes exhibit extreme high or low values in TA than RA (Fig. 3C). Not surprisingly, the log_2_(NP) *vs*. log_2_(RA) regression gave a slope of 1.04 – also significantly greater than 1 with a smaller than 1E-16 p-value, reflecting that more genes exhibit extreme high or low values in NP than RA (Fig. 4A). The trend is beyond experimental noise, as illustrated by the two RA biological replicates in figures 3A, 3C and 4A.

This sequential increase of selectivity is another architectural similarity of gene expression to the computer information retrieval process. As discussed earlier and shown in figure 1B, the sequential capacity decrease from HD to memory and to CPU cache in computers renders the information retrieval process sequentially more selective, *i.e*., focusing on smaller sets of information, necessitating efficient memory management (34).

### Quantification of cytoplasmic mRNA sequestration

The incorporation of the NP data enabled us to analyze another aspect of human gene expression – cytoplasmic mRNA sequestration. Upon nucleus export, not all mRNAs proceed directly to polysome association or degradation. Instead, many regulatory mechanisms selectively retain mRNAs. Sequestration has been used to refer to this phenomenon that mRNAs can remain stable, but not translating. We do not understand the roles of mRNA sequestration in cellular operation as well as those of transcription and translation.

To quantify individual mRNA sequestration levels, a mRNA sequestration index was calculated. We performed a log_2_(NP) ~ log_2_(RA)*log_2_(TA) linear regression, thus taking both RA and TA into consideration. The residue of this regression – the difference between observed log_2_(NP) value and expected value – quantifies raw mRNA cytoplasmic sequestration. It is well known that mRNA ORF length affects ribosome density on the mRNA and thus its distribution among the fractions in a cytoplasmic sucrose gradient fractionation experiment (57,58). As shown in figure 4C, our analysis also observed the effect of ORF length. The raw mRNA sequestration level negatively correlates with log_2_(ORF length), as illustrated by a loess regression. Thus, we adjusted the raw sequestration level with the loess regression, *i.e*., using the “residuals” component of loess regression output as our mRNA sequestration index, eliminating the ORF-length imposed complication in downstream functional analysis.

Given the overall architectural analogies from gene expression to computer information retrieval and the importance of memory-management in system operational efficiency in computers, we explored our dataset to investigate whether and how much the management principles would shed insights into cellular orchestration of transcriptome regulation and its entanglement with cellular operational efficiency. Specifically, we tested whether the principles also outline the relationship among TR, RA, TA and mRNA sequestration, thus bridging transcriptome regulation and cellular mitigation of the operational latency.

### The locality principles: spatial locality and transcriptional coregulation

First, we investigated the applicability of the spatiotemporal locality principles implemented in the OS kernel – the spatial locality for speculative HD-to-memory information loading and the temporal locality for information purging (35) (Fig. 1B) – in explaining common observations in gene expression analysis.

The spatial locality principle governs two inter-locked processes: HD information organization and HD-to-memory information loading (34,35) (Fig. 1B). Related information is stored spatially together to expedite data lookup and retrieval; concomitantly, the CPU tends to repeatedly use information from the same HD neighborhoods. Thus, upon a CPU memory request miss, *i.e*., the requested information is absent from the memory, the computer will speculatively load the whole neighborhood, instead of just the requested piece. Such speculative loading reduced the latency, as the information that the CPU will need soon is already pre-loaded into the memory. As the CPU will likely request the other information in the neighborhood soon, wasteful loading is also minimized; that is, the preloaded information will likely be used prior to having to be swapped out of the memory. In a word, the spatial locality principle strikes an optimal balance between the need for speculative loading prior to CPU request to minimize latency and the incurrence of wasteful loading. As the answer to the first question mark (?^1^) in figure 1B, this is obviously equivalent to co-transcriptional regulation of functionally related genes in the cells. In bacteria, the genes are often organized into poly-cistron operons, which is analogous to the organization of related information into a continuous string in a flat file in computers. In eukaryotic, though each gene is an independent transcription unit, functionally related genes often share common index, *e.g*., transcription factor binding sites; this scheme is analogous to the organization of related information into a computer database file in that each piece of information is directly addressable, but related pieces share common addressing indices.

### The locality principles: temporal locality and the mRNA stabilization-by-translation regulatory mechanism

The temporal locality principle governs the other half of memory management–identification of information already in the memory for purging to free up space for incoming information (34,35) (Fig. 1B). This principle takes advantage of the observation that the CPU request and usage tend to cluster at specific memory locations within a short period of time–hence the temporal locality. Thus, only memory locations not recently used by the CPU are identified for purging, to avoid having to quickly re-load them.

The cellular equivalent to memory information purging is mRNA degradation (Fig. 1B), and elaborate control of this process is a major factor in the “economics” of transcriptome regulation (59). Briefly, stabilization of an mRNA reduces the need for its transcriptional production and the associated energetic and metabolic overhead, as transcription consumes significant amount of ATP and nucleotide building blocks. It also enables the cell to bypass the time-consuming steps of transcription, processing and nucleus export for quick protein production. On the other hand, over stabilization of mRNAs generates added regulatory demands, for instance, to guard against aberrant protein production from the mRNAs, levying extra pressure on the regulatory machinery such as the RNA-binding proteins.

Stabilization-by-translation, a major mRNA degradation control mechanism (60), is functionally similar to the computer temporal locality principle, in that they both protect information that is actively requested/used at the execution level from being purged out of the intermediate step of the retrieval process. For instance, mRNAs with optimized amino acid codon composition generally have higher translation efficiency and, thus, are more stable (61). One major underpinning of this mRNA stability control is the EIF4F complex, which integrates the 7-methylguanosine-cap structure at the 5’ end of mRNAs and the polyA tails at the 3’ end, recruits ribosomes, and prevents exonuclease-mediated mRNA decay (62,63). Stabilization of actively translating mRNAs is intuitively understandable, as an actively translating mRNA is likely producing mission critical proteins. Moreover, the genome-to-proteome trend of enhanced selectivity is likely intertwined with mRNA degradation, the causative regulatory force for the TR-RA discrepancy described in figures 3 and 4A.

Thus, to address the 2^nd^ question mark (?^2^) in figure 1B, we hypothesized this stable-when-translating regulatory mechanism as cellular equivalent to the temporal locality principle. Our dataset enabled calculation of, as previously described, mRNA stability and translation indices to test this hypothesis (6):

- First, the log_2_(RA/TR) log-ratio, the stability index, estimates the degradation activity (Figs. 1 B and C, red arrows). TR is used in this ratio as mRNA production rate; due to extensive coupling of RNA processing to transcription and that transcription is on average more than three folds slower than RNA processing (12,13,64), transcription is the rate-limiting step in mRNA production and TR closely correlates with mRNA production rate for most genes (13). Under our experimental condition, most mRNAs should be in steadystate condition and follow the equations below:

■ 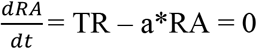 a: the degradation rate constant
■ TR = a*RA
■ a = RA/TR That is, log_2_(RA/TR) should estimate the degradation activity (Figs. 1 B and C, red arrows). Low degradation activity leads to higher RA than TR and thus positive log-ratio values, and high degradation activity the opposite, reflecting the increased selectivity from TR to RA due to mRNA degradation (Fig. 3A).
- Second, the log_2_(TA/RA) log-ratio is used, as commonly practiced, as mRNA translation index. High translation activity leads to higher TA than RA and thus positive log-ratio values, and low translation activity the opposite. The log-ratio reflects the steeper than 1 slope of the TA *vs*. RA regression (Fig. 3C), *i.e*., the increased selectivity from RA to TA, due to translation regulation.

To test the hypothesis, we first computed predicted RA (RA^Pred^) and TA (TA^Pred^) values by the respective log_2_(RA) *vs*. log_2_(TR) (Fig. 3A) and log_2_(TA) *vs*. log_2_(RA) (Fig. 3C) regressions, *i.e*., the “fitted.values” component of the outputs (or the corresponding points on the regression lines). The regression lines masked out the scatteredness of the data points due to the function-specific patterns of gene expression regulation and experimental noise, thus better reflecting the TR-to-RA and RA-to-TA increases in gene expression selectivity. We then computed the log_2_(RA^Pred^/TR) and log_2_(TA^Pred^/RA) log-ratios. If our hypothesis is correct, a positive correlation between the two was expected. This is, indeed, the case. As shown in a scatter plot in figure 5A, the two log-ratios have a correlation coefficient of 0.7, with a p-value lower than 1E-16, supporting mRNA stabilization-by-translation as the answer to the second question mark (?^2^) in figure 1B.

**Figure 5.**
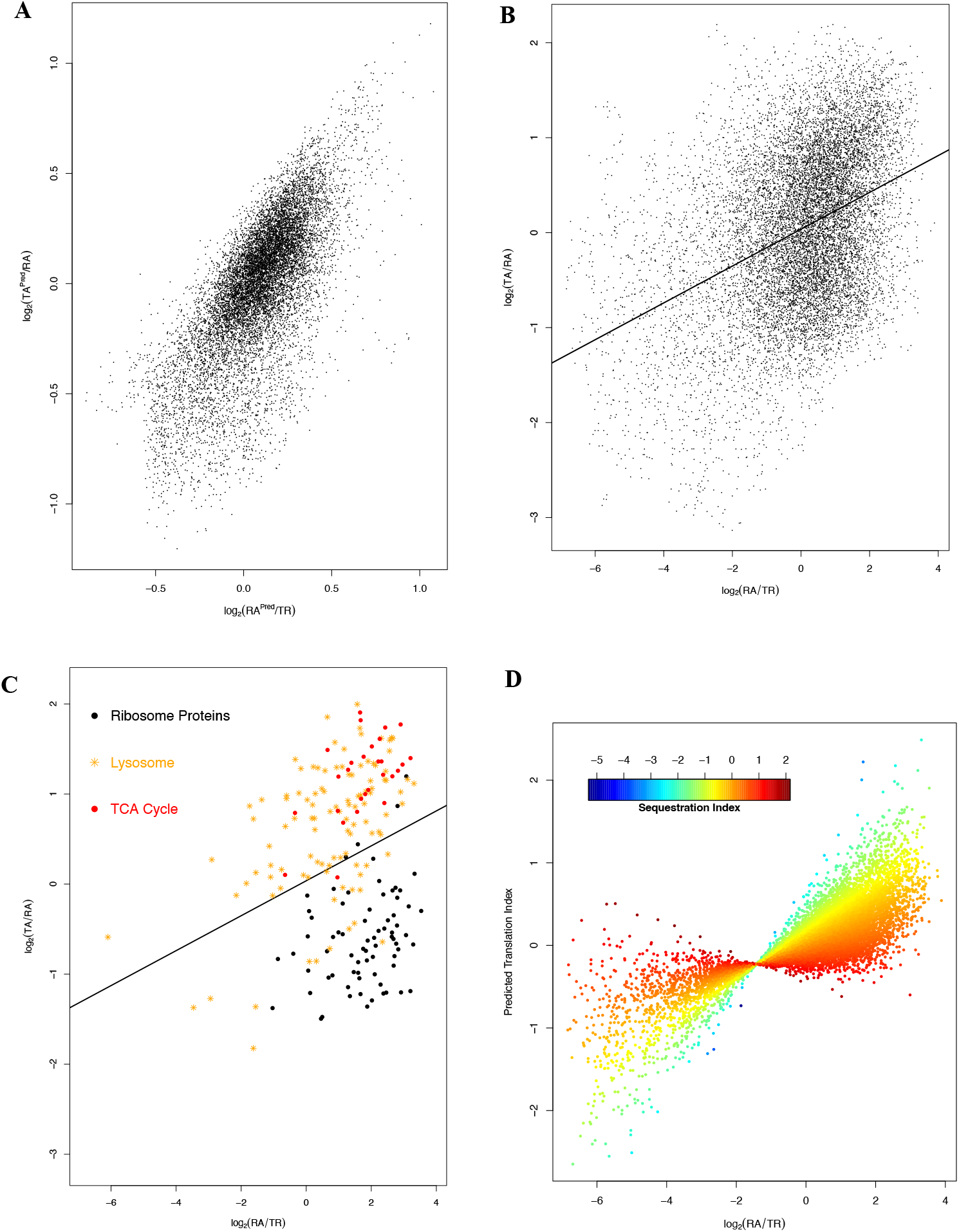
Analyzing the correlation between mRNA stability index and translation index. In **A**, the two indices were calculated with predicted RA and TA values by the linear regressions in figures 4A and 4C, respectively, *i.e*., log_2_(RA^Pred^/TR) and log_2_(TA^Pred^/RA). **B**: scatter plot of the two indices calculated with raw experimental RA and TA values, *i.e*., log_2_(RA/TR) and log_2_(TA/RA), exhibiting weaker correlation. The linear regression line is also shown. The positive correlation in A and B supports mRNA stabilization-by-translation as the answer to the second question mark (?^2^) in figure 1B. **C**: a scatter plot for the three KEGG functional groups as in figures 3B, 3D, 4B and 4D. The linear regression line from B is also shown. **D**: scatter plot of mRNA translation index predicted by a linear regression (log_2_(TA/RA) vs log_2_(RA/TR)*(sequestration index)) and the mRNA stability index (log_2_(RA/TR)). The data points are color-coded by the mRNA sequestration index.

Nevertheless, masking out the expected function-specific pattern of gene expression regulation is a significant drawback. Thus, a scatter plot of log_2_(TA/RA) *vs*. log_2_(RA/TR) is also shown (Fig. 5B). While positive correlation between the two log-ratios still exists as previously shown, the data points became more scattered. The correlation coefficient decreases to 0.39, reflecting the effect of function-specific mode of transcriptome regulation. The function-specific pattern was illustrated with the mRNAs for proteins in the TCA cycle, the lysosome and the ribosome (Figs. 3B, 3D and 5C). The three mRNA groups all exhibited overall higher-than-expected log_2_(RA) levels (Fig. 3B), but the ribosome mRNAs exhibited lower-than-expected TA levels (Fig. 3D). Consistently, ribosome mRNAs showed lower-than-expected log_2_(TA/RA) values, and lower log_2_(TA/RA)-log_2_(RA/TR) correlation than the lysosome and the TCA-cycle mRNAs (Fig. 5C). In short, the ribosome mRNAs defy the stabilization-by-translation regulatory mechanism, *i.e*., the temporal locality principle, while the TCA cycle and the lysosome mRNAs follow it.

Next, we performed a transcriptome-wide delineation of the function-specific pattern in conjunction with a further examination of the cell-computer analogy, searching for functional and mechanistic underpinning for the defiance of the mRNA stabilization-by-translation regulatory mechanism and thus the modesty of the log_2_(TA/RA)-log_2_(RA/TR) correlation.

### Defiance of the temporal locality principle: permanent caching of OS kernel in computers and function-specific cytoplasmic mRNA sequestration in the cells

As discussed earlier, computer software is organized into the kernel and the user programs (Fig. 1B). In computers with monolithic kernels, there is a clear boundary between the two. In computers with micro- or hybrid-kernels, though, the boundary becomes a bit blurred.

Concomitantly, there is another aspect of memory management – the differential management of the kernel and the user programs (34). User programs, as exemplified by internet browsers and Microsoft Word, are developed for special purposes, not allowed to access kernel memory area and controlled by the spatiotemporal locality principles. To the contrary, the kernel manages all the resources of the computer, *e.g*., CPU time, the memory and the hard drive. Instead of serving individual user programs, the kernel ensures a robust running environment for all of them; individual user program spontaneously interacts with the kernel indirectly, *e.g*., via OS system calls, to either request services for its own running needs or respond to kernel demands. Upon a kernel service request, a memory miss is operationally much more detrimental, potentially crashing the whole system. Thus, the kernel is permanently cached in the designated kernel memory area, as opposed to controlled by the locality principles, to ensure constant availability in a running computer.

We investigated whether and how the cells similarly organize cellular functions, and differentially regulate mRNAs for specific functional domains and those for core cellular functions that are critical for all cellular processes; that is, to answer the 3^rd^ question mark (?^3^) in figure 1B.

The moderate log_2_(TA/RA)-log_2_(RA/TR) correlation (Fig. 5B) suggests that, like the defiance of the locality principle by the OS kernel in computers, many mRNAs defy the stabilization-by-translation regulatory mechanism, as exemplified by the ribosome protein mRNAs (Fig. 5C). As discussed earlier, many other post-transcription regulatory mechanisms control mRNA stability in such manners, in that they stabilize mRNAs independent of, sometimes even accompanied by inhibiting, mRNA translation activity. That is, they maintain caches of mRNAs that are sequestered away from translation. One type of such regulators is the RNA-binding proteins (RBP); many of the more than 400 RBPs inhibit translation while the inhibited mRNAs remain stable, *e.g*., the regulation of insulin-like growth factor 2 (IGF2), beta-actin (ACTB) and beta-transducin repeat-containing protein (BTRC) mRNAs by insulin like growth factor 2 mRNA binding protein 1 (IGF2BP1) (65). Additionally, the microRNA (miRNA) regulatory system suppresses the translation of their target mRNAs and destines the mRNAs for sequestration in the p-body and/or the GW-body (66). While some target mRNAs are de-capped and degraded, a significant portion remains stable. The sequestration regulatory actions might occur at specific dedicated sub-cellular foci. For instance, the GW-body stores translationally inactive mRNAs (67). The P-body is also a repository for such mRNAs, though a portion of stored mRNAs is degraded. Another example is the cellular formation of stress granules upon exposure to environmental stressors. Generally, these regulatory mechanisms act through cognate binding sites in the mRNA 3’- or 5’-untranslated regions (UTR). As described earlier, their prevalence and importance in human cells are underscored by the observation that, on average, ~50% of a human mRNA are UTRs (6).

The caching principle in computer and the mRNA cytoplasmic sequestration are strikingly similar, in that they both retain selective information in the intermediate step regardless of their request/usage by the execution/decoding step. Thus, we explored our NP data and the mRNA sequestration index to investigate the applicability of the caching principle, *i.e*., explaining the mRNA sequestration pattern and the modesty of the log_2_(TA/RA)-log_2_(RA/TR) correlation.

Encouraging pattern was already observed with ribosome protein, TCA cycle and lysosome protein mRNAs. The ribosomal protein mRNAs exhibit higher NP relative to RA values (Fig. 4B), which is consistent with their low TA values (Fig. 3D) and defiance of the stabilization-by-translation regulatory mechanism. To the contrary, the TCA cycle and lysosome protein mRNAs exhibit largely lower NP relative to RA values (Fig. 4B), which is consistent with their high TA values (Fig. 3D). As shown in figure 4D, ribosome protein mRNAs thus exhibited higher sequestration levels than the TCA cycle and lysosome mRNAs. Not surprisingly, they seem defy the stabilization-by-translation regulatory mechanism, while TCA and lysosome protein mRNAs follow the regulatory mechanism (Fig. 5C). The pattern supports the notion that cytoplasmic sequestration level reflects whether the mRNA is controlled by the stabilization-by-translation regulatory mechanism.

Thus, the sequestration index was used to determine whether the function-specific mRNA sequestration illustrated in figure 4D is a genome wide pattern, that is, whether a pair of functionally related genes share similar mRNA sequestration levels. We calculated pair-wise difference in the sequestration indices (index^i^ – index^j^), termed sequ^diff^, among the genes. This sequ^diff^ parameter was then analyzed in relation to two measurements of functional relatedness of the gene pairs. Firstly, pair-wise Gene Ontology (GO) fingerprint similarity was used as measurement of the functional relatedness (see Materials and Methods for detail); the distribution of sequ^diff^ of gene pairs with different levels of GO similarities was compared. The sequ^diff^ should follow normal distributions with a mean of 0 and variances given by the following equation:

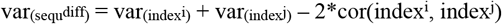

If the function specific pattern exists, the index^i^-index^j^ correlation should increase as GO similarity increases. Thus, the sequ^diff^ distribution should exhibit decreasing variance, *i.e*., the level of dispersion (see Materials and Methods for detail). As shown in figure 6A, this is indeed the case. The box-plots all indicated the same mean value of 0, but exhibited steadily decreasing levels of dispersion, *i.e*., lower and lower proportions of gene pairs have significantly non-zero sequ^diff^ values, along with increasing GO similarities. To display the trend quantitatively, the gene pairs were ordered by the GO similarity score and binned, with a bin size of 10000 and a 50% overlap between adjacent bins. As shown by a scatter plot in figure 6B, as the median GO similarity score in the bins increases, the SD of sequ^diff^ in the bins decreases; the blue horizontal line indicates the SD for all gene pairs (0.847) (see Materials and Methods for detail), and the black line the SD for gene pairs with no significant GO similarity. When the GO similarity scores are lower than 50, the data-points displayed scatteredness, though the overall trend is obvious. Above the similarity score of 50, the scattering disappears. In the second analysis, we used the protein-protein interaction data. The same trend as that in figure 6A was observed. The distributions of sequ^diff^ exhibited lower dispersion for gene pairs with interacting proteins, and the dispersion decreased along with increasing interaction quality scores (Figure 6C). Thus, a systematic function-specific pattern of mRNA sequestration was observed. The pattern is further confirmed, to be described later, by analysis of the KEGG functional gene sets (Fig. 6D).

**Figure 6.**
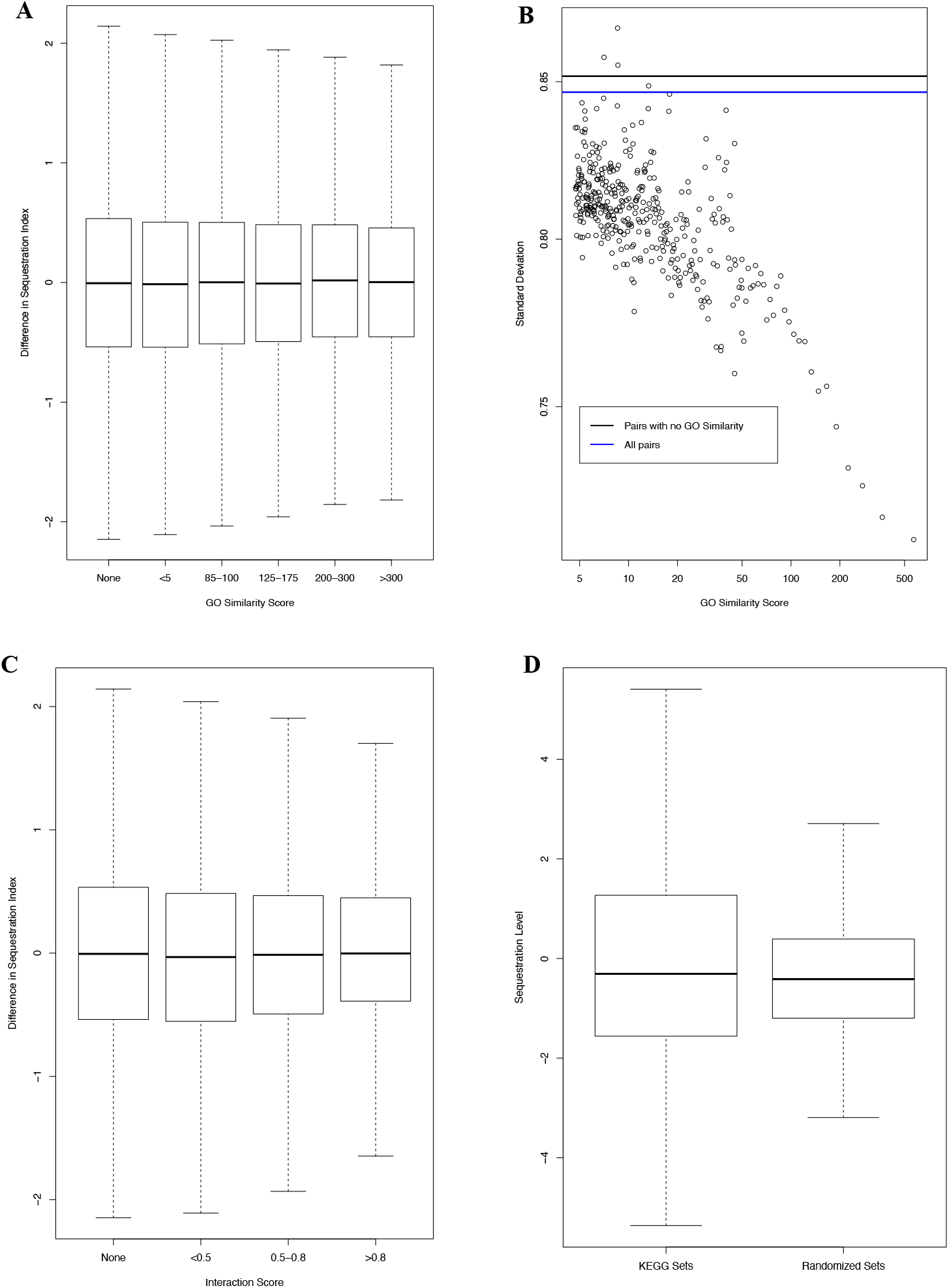
Function-specific mRNA sequestration pattern illustrated by genome-wide analyses. For each gene pair, their GO similarity and the differences between their sequestration index (termed sequ^diff^ here and in in text) were calculated. **A:** Comparative boxplot of the sequ^diff^ of gene pair sets with increasing levels of GO similarity. **B:** The gene pairs were binned by the GO similarity score. A scatter plot of the standard deviation (SD) of the sequ^diff^ versus the median GO similarity scores in the bins is shown; the black and blue horizontal lines denote the SDs for gene pairs with no GO similarity and all gene pairs, respectively. **C:** Comparative boxplot of the sequ^diff^ of gene pairs with varying protein-protein interaction scores. **D:** comparative boxplot of the sequestration levels of KEGG gene sets and randomized set, showing higher dispersion level of the KEGG sets. The randomized set has the same number of groups and group sizes.

Next, we tested whether high mRNA cytoplasmic sequestration levels are associated with the defiance of the stable-when-translating regulatory mechanism. The genes were ordered by their sequestration index. A sliding window analysis of the log_2_(TA/RA)-log_2_(RA/TR) correlation coefficient versus the sequestration index is shown in figure 7A, displaying a clear negative correlation. When the sequestration index is low, the genes have high log_2_(TA/RA)-log_2_(RA/TR) correlation, obeying the stable-when-translating regulation and thus the temporal locality principle. As the sequestration index increases, the log_2_(TA/RA)-log_2_(RA/TR) correlation steadily decreases. And mRNA cytoplasmic sequestration, thus the caching principle, progressively becomes the dominant regulatory mechanism. This transition is also shown schematically via a linear regression (log_2_(TA/RA) vs log_2_(RA/TR)*(sequestration index)) (Fig. 4D). Translation index values predicted by this regression was plotted versus the log_2_(RA/TR) log-ratio, with the data points color-coded by the sequestration index (Fig. 4D). At low mRNA sequestration levels, the data points display a clear positive correlation. As sequestration level increases, the correlation gradually disappears. Additionally, this regression effectively adjusted the log_2_(RA/TR) index with the sequestration index; not surprisingly, the adjustment increased correlation coefficient with the log_2_(TA/RA) index, moderately but statistically significantly, from 0.39 to 0.47. These results support cytoplasmic mRNA sequestration as cellular equivalent to the OS kernel caching in computers and the mechanistic underpinning for the defiance of the stable-when-translating regulatory mechanism.

**Figure 7.**
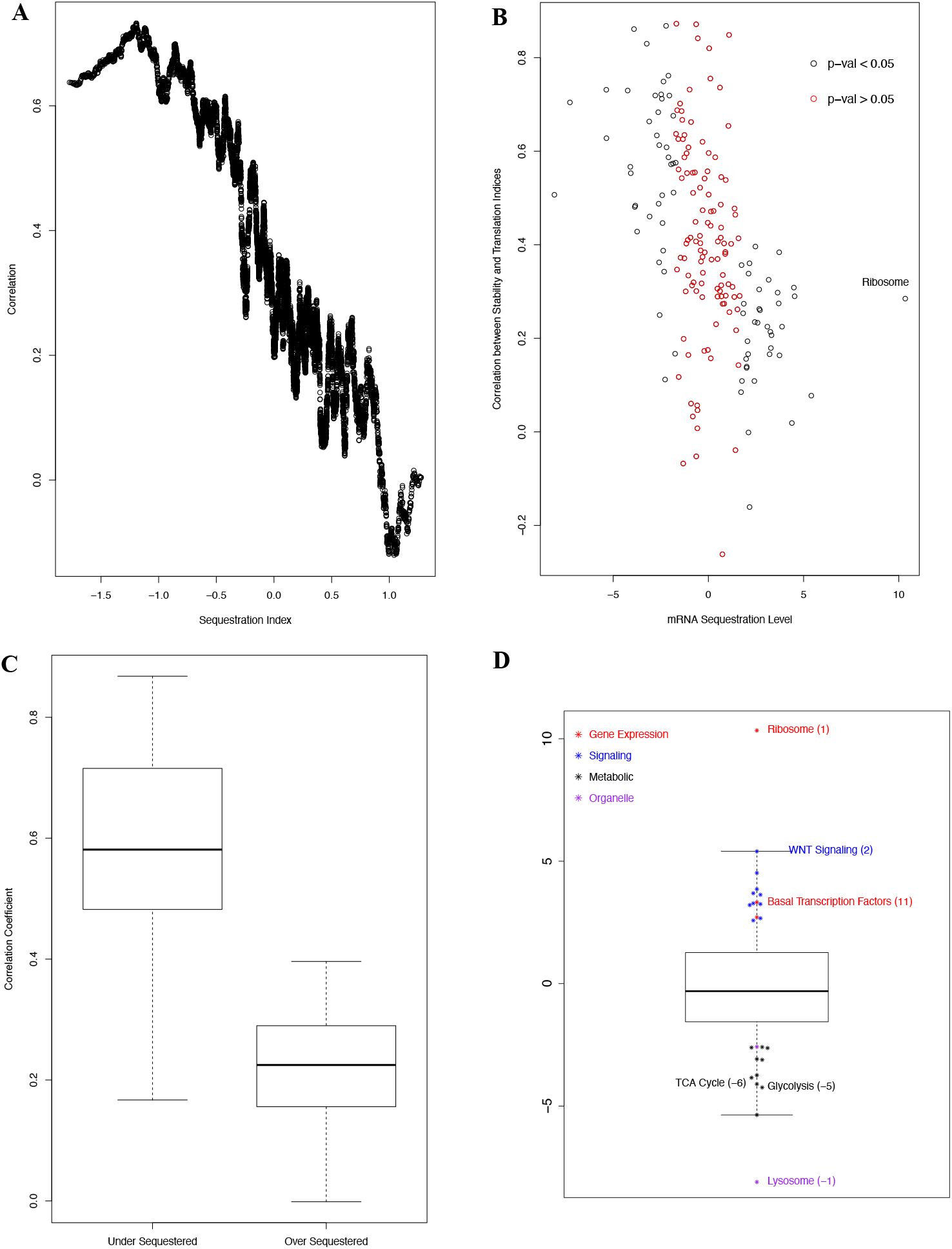
Negative relationship between mRNA sequestration index and the mRNA translation-stability correlation, as well as the function-specific pattern among the KEGG functional gene sets. **A:** Sliding window analysis of the relationship between mRNA sequestration index and the mRNA translation-stability correlation. **B:** Association of mRNA sequestration level and mRNA translation-stability correlation among the KEGG functional gene sets. For each gene set, mRNA sequestration level (in the form of t-scores) and translation-stability correlation coefficient were calculated. A scatter plot of the correlation coefficients versus mRNA sequestration levels is shown. The red data points denote groups with statistically insignificant sequestration levels (p-value > 0.05). The ribosome protein is labeled to illustrate the extreme sequestration level. **C:** comparative scatter plot of the correlation coefficients of under- and oversequestrated KEGG functional groups. Together with fig. 5D, the results support mRNA-sequestration *vs* memory-caching equivalence, answering the ?^3^ in fig. 1B. **D:** Over-sequestration of mRNA for gene-expression- and signaling-related KEGG gene sets in contrast to undersequestration for metabolism- and subcellular-organelle-related sets. Gene sets ranked within the top-20 most or least sequestered are shown. The data points are super-imposed onto a boxplot of the sequestration levels of the KEGG gene-set collection. Selective data points are labeled with gene set names and parenthesized ranks, with negative numbers denoting least sequestration ranking. Please see table 2 for full list of gene sets and ranking.

### Control of the information channel by equivalent mechanisms: caching of OS kernel in computers and cytoplasmic sequestration of mRNAs for gene expression machinery and their regulators in the cells

Our dataset provided an opportunity to identify cellular functions that are controlled by the cytoplasmic mRNA sequestration mechanism. Whether this mechanism regulates cellular functional equivalent to the computer OS kernel could then be determined (Fig. 1B, ?^3^). We downloaded the curated KEGG functional gene sets. Not surprisingly, ANOVA analysis (the sequestration-index versus the gene-sets) confirmed function-specific mRNA sequestration (F_185,8504_= 5.88, p-value < 2.2e-16); that is, the index values diverge across, but converge within, the gene sets. We then calculated gene-set level sequestration (in the form of t-scores), enabling schematic illustration of the function-specific pattern with comparative boxplots in figure 6D. The t-score distribution of the KEGG gene-sets exhibited significantly higher level of dispersion than that of same-sized sets of randomly selected genes; without the function-specific pattern, the dispersion should be similar. We also calculated the log_2_(TA/RA) *vs*. log_2_(RA/TR) correlation for each group. As shown in figure 7B, a negative correlation between the sequestration level and the log_2_(TA/RA)-log_2_(RA/TR) correlation was observed. Significantly over sequestered groups exhibited lower correlation than under sequestered groups (Fig. 7C).

Table 2 lists the respective top-20-ranked functional groups with highest- or lowest-sequestration levels. They are also shown schematically in Figure 7D. Metabolic pathways dominate undersequestrated groups; 9 of the top 20 are metabolic functions, with TCA cycle and glycolysis ranked at the 6^th^ and 5^th^, respectively. Some sub-cellular organelles are also under-sequestered; lysosome was ranked the 1^st^, and proteasome the 20^th^ (Figure 7D and Table 2).

**Table 2:** Top 20 ranked KEGG functions with least or most mRNA sequestration. Gene expression related terms are in red text, cellular signaling terms in blue, metabolism terms in green and subcellular organelle terms in purple text.

| Rank | Least Sequestered | Most Sequestered |
| --- | --- | --- |
| 1 | lysosome | ribosome |
| 2 | ecm_receptor_interaction | wnt_signaling_pathway |
| 3 | other_glycan_degradation | b_cell_receptor_signaling_pathway |
| 4 | cell_adhesion_molecules_cams | renal_cell_carcinoma |
| 5 | glycolysis_gluconeogenesis | long_term_potentiation |
| 6 | citrate_cycle_tca_cycle | mapk_signaling_pathway |
| 7 | dna_replication | chronic_myeloid_leukemia |
| 8 | asthma | colorectal_cancer |
| 9 | antigen_processing_and_presentation | neurotrophin_signaling_pathway |
| 10 | propanoate_metabolism | t_cell_receptor_signaling_pathway |
| 11 | valine_leucine_and_isoleucine_degradation | basal_transcription_factors |
| 12 | autoimmune_thyroid_disease | jak_stat_signaling_pathway |
| 13 | sphingolipid_metabolism | chemokine_signaling_pathway |
| 14 | galactose_metabolism | erbb_signaling_pathway |
| 15 | graft_versus_host_disease | acute_myeloid_leukemia |
| 16 | allograft_rejection | long_term_depression |
| 17 | steroid_biosynthesis | ubiquitin_mediated_proteolysis |
| 18 | o_glycan_biosynthesis | mtor_signaling_pathway |
| 19 | pyruvate_metabolism | pancreatic_cancer |
| 20 | proteasome | phosphatidylinositol_signaling_system |

On the other hand, the gene expression machineries and cellular signaling dominate oversequestrated functions (Figure 7D and Table 2). Ribosome and basal transcription factors were ranked at the 1 ^st^ and 11 ^th^, respectively; ubiquitin mediated proteolysis was ranked at the 17^th^. 10 of the top 20 functions are cellular signaling related; the WNT, MAPK, JAK-STAT and mTOR signaling were ranked at the 2^nd^, 6^th^, 12^th^ and 18^th^.

Thus, the gene expression machinery and their regulators in the cells and the OS kernel in the computers are not only functionally equivalent to each other, but also controlled by equivalent mechanisms. They act as the information flow channel in the respective systems. Similar to OS kernel serving core system functions in computers, the gene expression machinery and their regulators control essentially all cellular pathways by dynamically adjusting corresponding protein abundance. The gene expression machinery and their regulators are regulated by cytoplasmic mRNA sequestration – the cellular equivalent to memory caching of the OS kernel in computers. And it is likely not a coincident that the cache-controller is hardware-implemented, and the ribosome protein mRNAs are extremely sequestered (Fig. 7B); the two are functionally equivalent to each other, performing the extremely crucial step of information retrieval from the intermediate step to the execution/decoding level.

### Gene Ontology (GO) analyses further support the control of information channel by cytoplasmic mRNA sequestration

Encouraged by these observations, we investigated the function-specific pattern more comprehensively by performing the same analysis with the three GO gene sets: GO molecular function (MF), GO cellular component (CC) and GO biological process (BP). The GO BP set is much larger than other sets and contains many small sets, with as few as only three genes. We noticed that, for sets with 6 or less genes, their calculated sequestration levels exhibit high variance and are less reliable. Thus, we created a GO BP subset (GO BP (> 6)) that excludes these BP terms. The GO CC and MF sets did not have this problem. Table S1 lists the top 30 CC, MF, BP (>6) and BP terms with highest mRNA sequestration levels. Terms for gene expression machinery, their regulators and cellular signaling are color-coded (Table S1); the CC, MF and BP (>6) terms are also shown schematically in figure 8A. In table S2, all KEGG, GO CC, GO MF, GO BP (>6) and GO BP terms were listed in respective descending order.

**Figure 8.**
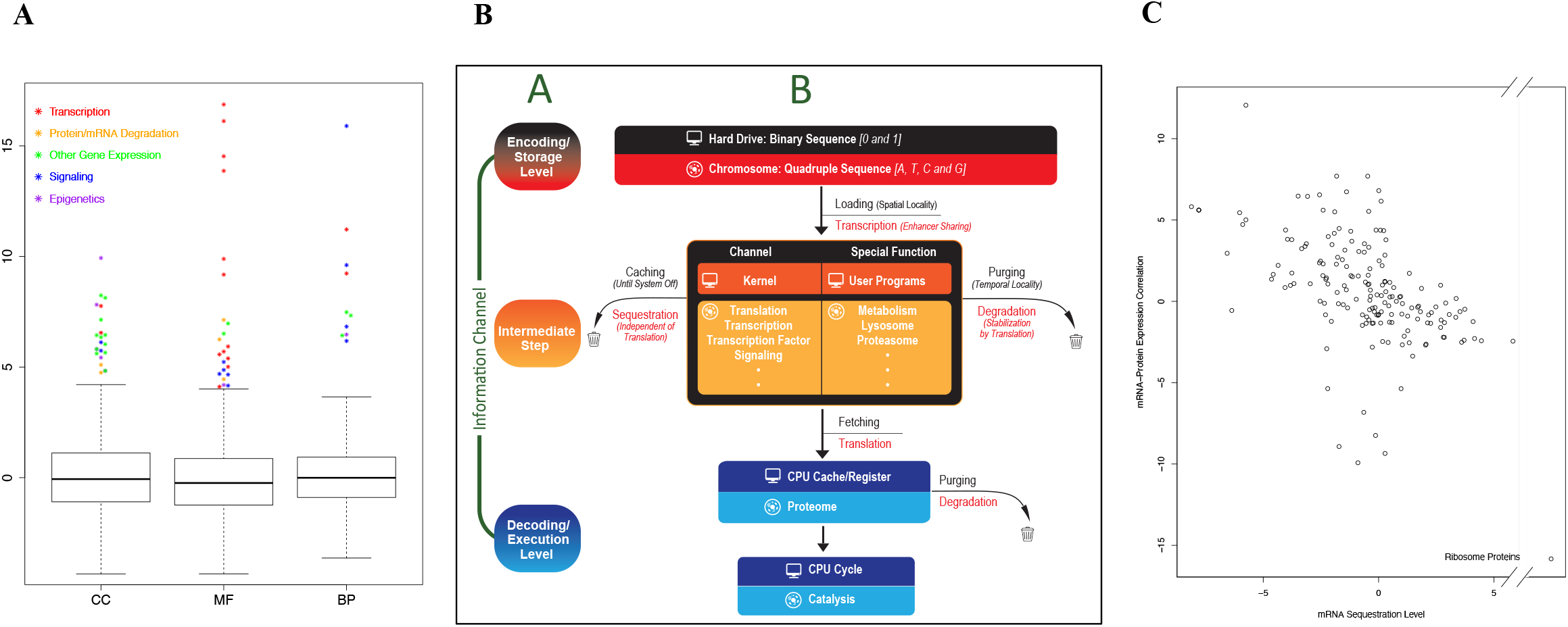
High mRNA-sequestration levels for gene expression and intracellular signaling (A), architectural and regulatory similarities between gene expression and computer information retrieval (B) and association between mRNA-sequestration and mRNA-protein expression discrepancy (C). **A:** Sequestration of mRNA for gene-expression- and signaling-related GO (CC, MF and BP) gene sets. Top-30 most sequestered gene sets are shown. Each data point is super-imposed, as in fig. 7D, onto the boxplot of the corresponding GO gene-set collection. Please see table S1 for the full list of gene names. Together with fig. 7D and tables 2 and S2, these results support mRNA sequestration for cellular equivalent to the permanently memory-cached computer operating system (OS) kernel (basal gene expression machinery, their regulators and intracellular signaling), answering the question mark (?^3^) in figure 1B. **B:** Improved analogy, upon that in fig. 1B, between cellular and computer instances of the information-theory. Sub-panel A copies the information-theory outline from fig. 1A. Sub-panel B depicts computer *vs* cell analogies as answers for the question marks (?^1,2,3^) in fig. 1B: spatial-locality *vs* enhancer-sharing for ?^1^; temporal-locality *vs* mRNA stabilization-by-translation for ?^2^ (see figs. 5A and 5B); and the memory-caching *vs* mRNA-cytoplasmic-sequestration analogy (see figs. 7A, 7B, 7C and 5D), and their control of the information channel (core system functions) (see figs. 7D and 8A), for ?^3^. The storage-to-execution textbox-width decrease denotes sequential capacity decrease, and increase in selectivity, in the systems (see Fig. 2). **C:** Association between mRNA-sequestration and mRNA-protein expression discrepancy. The correlation between mRNA and protein expression levels across the tumor samples is plotted *versus* mRNA sequestration level, one data point per KEGG gene-set, with ribosome proteins labeled to illustrate its extreme values.

As the KEGG functional group typically comprises of genes for different categories of biochemical activities, molecule-function-specific patterns might get lost in the above analysis. Indeed, the analysis of GO MF gene sets revealed such cases, and provided further evidence for cytoplasmic sequestration of mRNAs for gene expression machineries and their regulators. Transcription regulation and cellular signaling functions dominate the top 30 GO MF terms (Fig. 8A and Table S1). Most significantly, 12 of the 30 terms are related to transcription regulation; the top 4 ranked terms are all transcription factor (TF) related (Fig. 8A and Table S1, red text/symbol). Instead of forming their own biological process/pathway module, these transcriptional regulators often participate in different biochemical or signaling pathways. Thus, they are missed in our analysis of the KEGG biological process gene sets. Additionally, 5 of the 30 terms (ranked at 18^th^, 22^nd^ - 24^th^, and 28^st^) are related to intracellular signaling (Fig. 8A and Table S1, blue text/symbol). On the other hand, major metabolic enzymatic activities, such as lactate dehydrogenase, are ranked among the least sequestered terms (Table S2).

The GO CC and BP gene set analysis further confirmed over-sequestration of mRNAs for component of the information flow channel, as categorized below:

- As expected, many transcription regulation terms are also among the GO CC and BP terms (Fig. 8A and Table S1, red text/symbol).
- The transcription and the translation machineries, RNA binding proteins (RBP), translation regulators and protein transportation (Fig. 8A and Table S1, green text/symbol). In addition to the transcription factors/regulators, other components of the gene expression process are also enriched. Notably, 2 of the top 4 GO CC terms (2^nd^, 3^rd^) are all ribosome related; ranked at the 12^th^ is the mRNA-RBP complex. The term “nucleolus” and “fibrillar_center” are related to ribosome biogenesis and ranked at the 7th and 27^th^, respectively. The “nuclear _speckle”, the sub-nuclear structure enriched of proteins involved in transcription and mRNA splicing, is ranked at the 17^th^. As for GO MF terms, translation termination is ranked at the 11 ^th^. And three of the GO BP terms (ranked at 17^th^, 19^th^ and 26^th^) are related to translation regulation and protein transportation (Fig. 8A and Table S1, green text/symbol).
- The mRNA and the protein degradation machineries. Proteolysis and mRNA degradation are equivalent to information purging out of the CPU cache and the memory, respectively. Consistently, “ubiquitin-mediated-proteolysis” is the 17^th^-ranked most sequestered KEGG term (Fig. 7D and Table 2). Not surprisingly, three related terms are ranked at the 9^th^, 12^th^ and 26^th^ among GO MF terms (Fig. 8A and Table S1, orange text/symbol). Additionally, the GO CC gene set analysis revealed over-sequestration (ranked at the 24^th^ and 29^th^) of mRNAs for two related terms (Fig. 8A and Table S1, orange text/symbol); and the term “CCR4_NOT_complex” is ranked at 45^th^ (Table S2).
- Intracellular signaling terms (Fig. 8A and Table S1, blue text/symbol). The GO CC terms “phosphatase_complex” and “PI3-kinase complex I” terms are ranked at the 14^th^ and 18^th^, respectively. And four such terms are ranked at the 3^rd^, 9^th^, 21^st^ and 29^th^ among the GO BP terms.
- To the contrary, and as expected, subcellular organelles, such as lysosome and endoplasmic reticulum (ER), are ranked among the least sequestered GO CC terms, and major metabolic processes among the least sequestered GO BP terms (Table S2).

Moreover, the analysis of GO gene sets also revealed significant enrichment of epigenomic terms. Chromatinization and further condensation of the genome imposed further gene expression time delay and, if not mitigated, severe system latency in eukaryotic cells. Proteins for the epigenomic functions are thus part of the genome-to-proteome information flow channel. It is therefore not surprising that their mRNAs are controlled by cytoplasmic sequestration as well (Fig. 8A and Table S1, purple text). The terms “chromatin”, “chromosome” and “PML_body” are the 1^st^, 4^th^ and 22^nd^ ranked most sequestered GO CC terms, respectively; the term “histone_h3_k36_trimethylation” is the 25^th^ ranked GO BP term; and “histone_acetyltransferase_activity_h3_k23_specific” is the 27^th^ ranked GO MF terms. Additionally, the terms “chromatin_organization”, “covalent_chromatin_modification” and “chromatin_remodeling” are all ranked within the top 200 (67^th^, 106^th^ and 191 ^st^) out of the more than 7000 GO BP terms (Table S2).

Thus, we observed evidence that the gene expression machinery and their regulators are equivalent to the computer OS kernel, and the equivalence is multi-faceted. Firstly, they are both part of the information body that dynamically flows from the storage level to the execution/decoding level. Secondly, they serve core system functions instead of specialized functions, acting as master controller of the dynamic information flow process. That is, as discussed earlier, they are the information channels in the respective system. Thirdly, concomitantly, they are similarly granted preferential status to ensure ready availability to the execution level. In running computers with monolithic kernel, the kernel permanently resides in memory, not subject to the temporal-locality purging policy. And the cache-controller is hardware-implemented due to its operational importance. In the cells, the cognate mRNAs are regulated by cytoplasmic sequestration, defying the stable-when-translating regulatory mechanism. Not surprisingly, mRNAs for the ribosome, the cellular equivalent to the computer cache-controller, are extremely sequestrated (Figs 4B, 4D and 7B). In short, mRNA sequestration acts as the equivalent to permanent caching of OS kernel in the computer, answering the 3^rd^ question mark (?^3^) in figure 1B.

### Cytoplasmic mRNA sequestration explains partially mRNA-protein expression discrepancy

We were intrigued by whether and how much mRNA sequestration contributes to mRNA-protein expression discrepancy, as sequestered mRNAs can be quickly released to form polysome to enable translation activity fluctuation without mRNA abundance changes. As discussed earlier, this discrepancy is frequently observed in proteogenomic studies (29–31). Fortunately, one proteogenomic study measured mRNA and protein abundance in a collection of colorectal tumor samples, from which the HCT116 cells was also derived. Thus, the proteogenomic datasets were analyzed in conjunction with our mRNA sequestration index and the KEGG gene sets. For each gene in the dataset, we calculated the correlation coefficient between its mRNA and protein expression levels across the tissue samples. For each KEGG gene set, we identified its member genes in the datasets, re-calculated the mRNA sequestration level (in the form of a t-score), and calculated protein-mRNA correlation level (in the form of a t-score between their protein-mRNA correlation coefficients and those of all the genes in the dataset). As shown in figure 8C, a negative correlation was observed; high mRNA sequestration is associated with poor protein-mRNA correlation. Once again, ribosome protein is the outlier, *i.e*., with extremely low protein-mRNA correlation.

## Discussion

The genome-to-proteome information flow process and its centrality to cellular operation are encapsulated in the molecule biology central dogma. However, how the cells mitigate the latency imposed by the tardiness of this process remains unknown, perhaps due to current practical treatment of gene expression and cellular operational efficiency as separate conceptual and research domains. And our understanding of the long-observed gene expression complexity remains fragmentary. We studied this process as an instance of the Shannon information theory, and hence the gene expression machinery and their regulators as the information flow channel. This abstraction prompted a multi-omic comparative analysis with the dynamic HD-memory-CPU information flow–the instance of the same theory in support of system operation in computer. As summarized in figure 8B, we observed further architectural similarity between the gene-mRNA-protein gene-expression and the HD-memory-CPU information-retrieval process in computer. The analysis revealed evidence that the optimization principles implemented in computers to mitigate the operational latency also act in the cells, outlining cellular orchestration of multiple transcriptome regulatory mechanisms and the relationship among TR, RA, TA and mRNA sequestration (Fig. 8B). Our results also have the power to partially explain the frequently observed mRNA-protein expression discrepancy (Fig. 8C). That is, this study closes the practical gap to enable a conceptual bridging of transcriptome regulation and cellular operational latency mitigation.

Transcriptome regulation in human cells exhibits inherent, prohibitively enormous, complexity. Conceivably, the complexity evolved hand-in-hand with the complexity of cellular architecture and processes from bacteria to eukaryotes and to metazoans such as humans. The subcellular components and their associated functions become increasingly compartmentalized or modularized, *e.g*., the formation of the nucleus and resultant spatial separation of translation from transcription. While the gene counts sequentially increase, chromatinization and increasing chromosome condensation render transcription activation and elongation progressively slower and costlier. Not surprisingly, the evolutionary emergence of a myriad of post-transcriptional regulatory mechanisms accompanies that of metazoans. The examples include the microRNA regulatory system and the sets of RNA binding proteins, and more are sure to be un-covered. Unfortunately, the high levels of complexity also render direct theoretical inquiry for the underpinning principles difficult, if not impossible.

Our cell-to-computer analogy based on the respective implementation of the information theory in support of system operations seems helpful. The analogy led to a multi-omic measurement of vital gene expression steps, aiming to test the explanatory power of computer latency-mitigation principles. The analysis revealed additional architectural similarity; the sequential genome-to-proteome enhancement of selectivity was observed (Figs. 2, 3 and 4 and Table 1), in parallel to the sequentially HD-to-CPU lower capacity, and thus higher selectivity, in the computer (Fig. 8C); and mRNA degradation is equivalent to computer memory information purging. We also revealed regulatory similarities. The genomic organization of the genes was re-interpreted in the spirit of the spatial locality principle, and we generated experimental evidence that mRNA stabilization-by-translation is analogous to the temporal locality principle that governs selective information purging from the memory. That is, the spatiotemporal locality principle and its critical role in computer latency mitigation facilitate an interpretation of genomic organization and co-transcription regulation, as well as mRNA stability control by translation activity, in the context of cellular operational efficiency (Fig. 8C).

Moreover, our dataset provided an opportunity to tackle another transcriptome regulation complexity – the defiance of the locality principle. As discussed above, many other mechanisms regulate mRNAs, often resulting in mRNA sequestration/caching, *i.e*., stabilization of translationally inactive mRNAs. Sequestration/Caching analysis mandates simultaneous monitoring of mRNA stability and translation activity. The stability data cannot tell, by itself, whether the stabilization is due to translation activity or the caching mechanisms. Similarly, the translation activity data alone is powerless in assessing mRNA stability. Our transcriptome-wide depiction of the relationship between mRNA stability and translation activity met the prerequisite. This relationship, along with the public functional annotation of the genes, provided a framework for assessing mRNA sequestration and determining the mode of regulation of individual cellular functions, *i.e*., the extent to which their mRNAs are controlled by either caching or translation activity. Indeed, our analysis of cytoplasmic non-polysome-associated mRNAs confirmed that highly sequestered mRNA defies the locality/stable-when-translating regulatory mechanism. We observed that mRNAs for the gene expression machinery are highly cached, and so are mRNAs for their regulators and intracellular cellular signaling proteins. On the other hand, mRNAs for many metabolic pathways, such as glycolysis and TCA cycle, are much under-cached, and so are the mRNAs for structural proteins of some subcellular organelles, such as the lysosome and proteasome.

This mRNA sequestration pattern reinforced the cell-to-computer analogy (32,55), with a notion that differential control of core system functions and specialized functions occurs in a similar manner in the two systems (Fig. 8C). The gene expression machinery, their regulators, and the signaling molecules constitute the information flow channels in the cells; the basic machinery carries out the flow, and their regulators and cellular signaling determine specificity, *i.e*., dynamic allocation of the channel capacity among biochemical pathways. In computers, the HD-to-CPU information flow channel is arguably the OS kernel; the well-defined kernel functions (file system, memory management, and process control/scheduling) collectively accomplish the task. It is striking that, in both systems, channel components are themselves part of the flow process and given preferential status – mRNA sequestration and memory caching, respectively. While the similarity between computers and cells has been widely noticed and computers always have an OS, whether the cells have equivalent to the OS was questioned (55). Our results suggest that the gene expression machinery, regulators of the machinery, and intracellular signaling are arguably the answer. Thus, this study not only benefited from, but also further extended, the cell-to-computer analogy and the application of the information theory.

Our usage of computer optimization principles as a stepping-stone to tackle, indirectly, the complexity of the cellular operation is consistent with the tradition of using simpler model systems in biological research. For instance, the yeast *Saccharomyces cerevisiae* has long served as a model organism for understanding higher eukaryotes. Similarly, neural science and development biology have long used the worm *Caenorhabditis elegans* conveniently as the model. It is also consistent with an underlying notion in systems biology, that is, to explore similarities between biological and engineered complex systems (68).

To the best of our knowledge, this is the first report of the quantification of mRNA caching activity. And our results point to future investigation to further advance our understanding of this pervasive phenomenon. This analysis represents a snapshot of actively growing log-phase cells, which are not homogeneous cell populations. Instead, they are a mixture, for instance, of cells in different cell cycle stages. On the other hand, it is reasonable to assume that the mRNA caching activity is dynamic, *i.e*., cell cycle stage, cell type, and growth condition specific. Thus, there exists a need to apply our approach to multiple cell types and time-series experiments, during which the cells transition through physiological processes or adjust to stresses. This way, we will know when and why mRNA caching occurs. We will also know when the cell releases cached mRNAs into the polysomes for quick protein production. That is, we will be able to delineate the functional and operational advantages endowed to the cells by mRNA caching.

We would also like to point out that this analysis quantified the overall mRNA caching activity, not focusing on individual caching mechanisms or subcellular foci. On the other hand, we have learned a lot about specific caching mechanisms. For instance, the P-body has long been known as a repertoire for non-translating mRNAs. It was shown to act as sub-cellular foci for miRNA-mediated mRNA caching/sequestration (66). Some stored mRNAs get degraded, but some recycle back for active protein production. Recently, the GW-body was identified as another subcellular foci for translationally repressed mRNAs. And a large number of RBPs have been discovered in the human genome. Some of the RBPs are undoubtedly involved in mRNA caching (1,2). However, we believe there is a lot to be learned yet. Our results should be a good framework for the functional study of these subcellular foci and RBPs.

In summary, we described a mechanistic and functional study of the complexity of transcriptome regulation via a computer *versus* cell analogy (Figs. 1 and 8B). Human gene expression is metabolically expensive, *e.g*., consuming large amounts of building blocks as well as ATP and GTP as the energy source. The process is also time-consuming, imposing potential operational delays. Thus, there is a need to fine-tune the process for an optimal tradeoff among multiple cellular “economics” factors. Our analyses suggest the spatiotemporal locality and the caching principles are integral to such optimization schemes. The functional and regulatory equivalent between the storage-to-execution information flow channels of the cell and the computer is striking and begs for further investigation. They might be a pure coincidence of the biological evolution of the cells and the competition-driven evolution of computers in the market. Alternatively, they might be the manifestation of fundamental organizational and operational principles of complex systems. We will not be surprised, though, if the latter turns out to be the case.

